# Comparison of Prognostic Accuracy of the quick Sepsis-Related Organ Failure Assessment between Short- & Long-term Mortality in Patients Presenting Outside of The Intensive Care Unit – A Systematic Review & Meta-analysis

**DOI:** 10.1101/296459

**Authors:** Toh Leong Tan, Ying Jing Tang, Ling Jing Ching, Noraidatulakma Abdullah, Hui-min Neoh

## Abstract

**Objective:** In year 2016, quick Sepsis-Related Organ Failure Assessment (qSOFA) was introduced as a better sepsis screening tool compared to systemic inflammatory response syndrome (SIRS). The purpose of this systematic review and meta-analysis is to evaluate the ability of the qSOFA in predicting short- and long-term mortality among patients outside the intensive care unit setting.

**Method:** Studies reporting on the qSOFA and mortality from MEDLINE (published between 1946 and 15^th^ December 2017) and SCOPUS (published before 15^th^ December 2017). Hand-checking of the references of relevant articles was carried out. Studies were included if they involved inclusion of patients presenting to the ED; usage of Sepsis-3 definition with suspected infection; usage of qSOFA score for mortality prognostication; and written in English. Study details, patient demographics, qSOFA scores, short-term (<30 days) and long-term (≥30 days) mortality were extracted. Two reviewers conducted all reviews and data extraction independently.

**Results and Discussion:** A total of 39 studies met the selection criteria for full text review and only 36 studies were included. Data on qSOFA scores and mortality rate were extracted from 36 studies from 15 countries. The pooled odds ratio was 5.5 and 4.7 for short-term and long-term mortality respectively. The overall pooled sensitivity and specificity for the qSOFA was 48% and 85% for short-term mortality and 32% and 92% for long-term mortality, respectively. Studies reporting on short-term mortality were heterogeneous (Tau=24%, I^2^=94%, P<0.001), while long-term mortality studies were homogenous (Tau=0%, I^2^<0.001, P=0.52). The factors contributing to heterogeneity may be wide age group, various clinical settings, variation in the timing of qSOFA scoring, and broad range of clinical diagnosis and criteria. There was no publication bias for short-term mortality analysis.

**Conclusion:** qSOFA score showed a poor sensitivity but moderate specificity for both short and long-term mortality prediction in patients with suspected infection. qSOFA score may be a cost-effective tool for sepsis prognostication outside of the ICU setting.

## INTRODUCTION

Sepsis is a syndrome of uncertain pathophysiology that is recognized by a group of clinical signs and symptoms in patients with suspected infection ^1^. Sepsis is a significant cause of mortality worldwide. In the last decade, an estimated 31.5 million sepsis patients have been treated globally per year, including 5.3 million deaths due to sepsis ^2^. The diagnosis of sepsis is challenging, as a reliable test for its early confirmation is not available. Given the morbidity and mortality of sepsis, the ability to perform risk stratification in the early phase of a patient’s illness may help physicians to effectively manage and improve their outcome.

The Third International Consensus Definitions for Sepsis and Septic Shock (Sepsis-3) defined sepsis as life-threatening organ dysfunction caused by a dysregulated host response to infection^1^, known as severe sepsis in previous definitions ^3^. The formerly used Systemic Inflammatory Response Syndrome (SIRS) criteria for early identification of sepsis was considered impractical and inefficient ^4^. Subsequently, Sepsis-3 proposed using the quick Sepsis-Related Organ Failure Assessment (qSOFA) as a new risk stratification tool to distinguish patients who are likely to have sepsis under the new definition. The qSOFA criteria recommend screening patients for three clinical signs i.e. tachypnoea, altered mental status, and hypotension, rather than requiring blood tests. Ongoing efforts have been directed toward examining the ability of qSOFA to predict poor outcomes in patients with infection ^5^. However, several studies have suggested that qSOFA criteria lack accuracy for predicting mortality in patients outside the intensive care unit compared to other early scoring systems ^6,7^. Since the qSOFA is a new scoring system, the clinical practicality of these criteria as a screening tool for infectious patients has not been fully evaluated.

The intention of this systematic review and meta-analysis was to evaluate qSOFA as a mortality predictor in patients presenting outside of the intensive care unit (ICU). We hypothesized that qSOFA can predict short- and long-term mortality in sepsis patients. The prognostic accuracy of qSOFA score for both short- (≤30 days) and long-term (>30 days) mortality was analysed.

## METHODS

### Study Eligibility Criteria and Search Strategy

A systematic review and meta-analysis of the literature was conducted to identify relevant studies regarding the role of the qSOFA in mortality prognostication among patients with suspected infection presented outside of the intensive care unit after obtaining consent from UKM Research Ethical Committee (UKM PPI/111/8/JEP-2017-769). We used MEDLINE via Ovid Medline to conduct a comprehensive search of health science journals (published between 1946 and 15 December 2017) and SCOPUS (published before 15 December 2017); Hand-checking of the references of relevant articles was carried out. The search team comprised of three clinicians, a statistician and a scientist. The search strategy involved a combination of the following 2 sets of keywords 1) ‘quick sequential organ failure assessment’ OR ‘quick SOFA’ OR ‘qSOFA’ OR ‘quick sepsis related organ failure assessment’ and 2) ‘mortalit*’. This meta-analysis was registered in PROSPERO (CRD42017079364, http://www.crd.york.ac.uk/PROSPERO/display_record.php?ID=CRD42017079364). The search strategies were shown in S1 Table.

### Identification and Selection of Studies

Study selection was performed based on their titles or abstracts, and only studies which appeared to fulfil the eligibility criteria were selected for full-text review. To be included, studies must fulfil the following criteria: inclusion of patients presenting to the ED; usage of Sepsis-3 definition with suspected infection; usage of qSOFA score for mortality prognostication; and written in English. Papers were excluded if they were: related to review articles; articles without complete texts; or animal studies.

### Data Extraction and Study Appraisal

The selection of papers inclusion in this review was completed in four phases. First, an initial search of the selected databases was performed using the pre-specified keywords to identify relevant keywords and index terms. Second, a thorough search was conducted in which papers that failed to meet the inclusion criteria based solely on their titles and abstracts were excluded. In the third phase, the remaining papers from the second phase were extensively reviewed, and papers that did not meet our inclusion criteria were excluded. Finally, all relevant data from the included papers was subjected to meta-analysis to determine conclusions regarding the proposed hypothesis.

After the initial screening of titles and abstracts by two independent reviewers, who are clinicians, incomplete articles were removed. The remaining papers were screened again by the two reviewers. To minimize errors, both reviewers were trained under a consensus standard and then practiced using several articles for calibration. Any discrepancies were resolved through discussion with a third reviewer who is an Emergency Physician. We applied the QUADAS-2 (Quality Assessment of Diagnostic Accuracy Studies) criteria to assess the quality of all selected articles. The risk of bias of each included study was summarized (S2 and S3 Tables). Data extraction was conducted independently in a standardized manner with a data collection form. Study data including author, publication year, type of study conducted, brief description of the study population/sample and methods used in the study, index/reference time interval, and mortality outcome were extracted from the full text of each article and summarized in detail (S4 and S5 Tables). In-hospital mortality was categorized as short-term mortality. This analysis was reported according to Transparent Reporting of Systematic Reviews and Meta-Analyses (PRISMA) guideline. A flow diagram of study identification and articles selection for the meta-analyses can be found in Figure 1.

**Figure 1.**
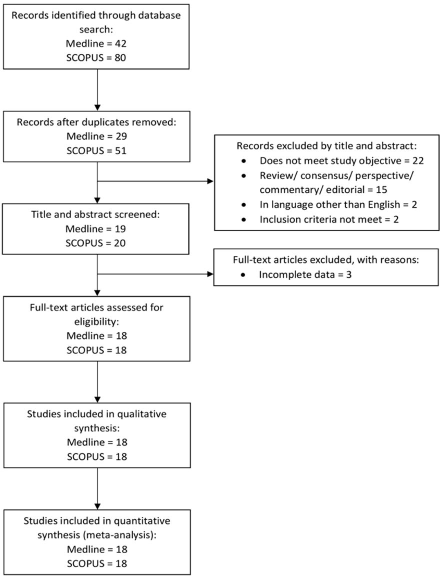
Identification and Selection of Articles for Meta-analysis. Flow chart shows process of article selection and exclusion throughout the study.

### Statistical Analysis

The primary statistical analysis was performed using the Review Manager 5 (Version 5.3.5) software manufactured by Cochrane Community and the Comprehensive Meta-Analysis Software (CMA, Version 3) manufactured by Biostat. Based on this model, the pooled sensitivity, specificity and the odds ratio (OR) with the 95% CI were determined. Random effects model was used to report short- and long-term mortality individually with estimates of sensitivity, specificity and ORs. The I^2^ statistic was calculated to determine the proportion of between-study variation caused by heterogeneity, with suggested thresholds for low (25%-49%), moderate (50%-74%), and high (75%) values. The publication bias of the included studies was assessed using effective sample-size funnel plot (OR values vs sample size of each study), Begg-adjusted rank correlation tests and the Egger regression asymmetry test for small study effects.

### Data availability

The authors declare that all data supporting the findings of this study are available within the paper and its supplementary information files.

## RESULTS

The search identified relevant studies from MEDLINE via Ovid Medline (1946 to 15 December 2017) and SCOPUS database (through 15 December 2017). The numbers of relevant records identified in MEDLINE and SCOPUS were 42 and 80, respectively, for a total of 122 references retrieved from the electronic database. Forty-two records were identified as duplicates and were removed from our selection. Of the 80 remaining references, 41 were excluded based on titles and abstracts: 22 did not meet the primary objective of our review, two did not meet our inclusion criteria, two studies were published in languages other than English, and 15 were other articles including review, consensus, perspective, commentary and editorial papers. The full texts of 39 studies were successfully retrieved. Three papers were excluded due to incomplete data in study (S6 and S7 Tables). The authors of the three studies were contacted through electronic mail, but no response were received. Finally, 36 studies fulfilled the inclusion criteria and were included. The characteristics of the included studies ^5–40^ are summarized in Table 1.

**Table 1.** Summary of Characteristics of Studies Included

| Source | No. of Participants | Mean Age, y | Men, No. (%) | Main Inclusion Criteria | Outcome |
| --- | --- | --- | --- | --- | --- |
| **Short-term mortality** |  |  |  |  |  |
| April (8), 2017 | 214 | 68 | 126 (59%) | ED patients admitted to any ICU with suspected or proven infection | In-hospital mortality |
| Askim (9), 2017 | 1535 | 61<sup>a</sup> | 813 (53%) | New onset of suspected or confirmed infection according to the ESS47 | 30-day mortality |
| Brabrand (10), 2016 | 3824 | 65<sup>a</sup> | 2426 (63%) | Patients presenting or discharged with suspected infection | In-hospital mortality and/or ICU stay >3days |
| Chen (11), 2016 | 1641 | 73<sup>a</sup> | 968 (59%) | Patients with CAP or healthcare-associated pneumonia | 28-day mortality |
| Churpek (6), 2017 | 30677 | 58 | 14561 (47%) | Patients with suspicion of infection in wards or ED | 28-day mortality |
| Churpek (12), 2017 | 53849 | 57 | 24719 (46%) | Patients meeting suspicion criteria in ED or wards | In-hospital mortality |
| DeGroot (13), 2017 | 2280 | 61 | 1315 (58%) | ED patients with suspected infection | In-hospital mortality |
| Donnelly (14), 2017 | 2593 | NA | NA | Admitted patients who meet SIRS criteria, SOFA and qSOFA criteria | 28-day mortality |
| Finkelsztein (15), 2017 | 152 | 64<sup>a</sup> | 83 (55%) | Patients with suspicion of infection admitted to the medical ICU from emergency department or hospital wards | In-hospital mortality |
| Forward (16), 2017 | 161 | 70 | 89 (55%) | Non-ICU inpatients who triggered the hospital SK pathway with acute deterioration and suspected or proven infection | In-hospital mortality |
| Freund (17), 2017 | 879 | 67<sup>a</sup> | 465 (53%) | Patients admitted to ED with clinical suspicion of infection | In-hospital mortality |
| Giamarellos (18), 2017 | 3436 | NA | NA | Patients with signs of infection | 28-day mortality |
| Gonzalez (19), 2017 | 1071 | 84 | 544 (51%) | Patients ≥75 years old clinically diagnosed with acute infection in ED | 30-day mortality |
| Haydar (20), 2017 | 199 | 71<sup>a</sup> | 109 (55%) | ED patients treated for suspected sepsis | In-hospital mortality |
| Henning (21), 2016 | 7637 | 60 | 3799 (50%) | ED patients admitted to the hospital with an infection-related diagnosis | In-hospital mortality |
| Huson (22), 2017 | 329 | 34<sup>a</sup> | 125 (38%) | Patients with suspected infection with ≥2 SIRS criteria | In-hospital mortality |
| Huson (23), 2017 | 458 | 35<sup>a</sup> | 243 (53%) | patients admitted to the adult medical ward with suspected infection | In-hospital mortality |
| Hwang (24), 2017 | 1395 | 65<sup>a</sup> | 787 (56%) | Patients who received a diagnosis of severe sepsis or septic shock during ED stay | 28-day mortality |
| Kim (25), 2017 | 615 | 54 | 204 (33%) | Patients with fever and chemotherapy-induced neutropenia | 28-day mortality |
| Kim (26), 2017 | 125 | 76 | 78 (62%) | Patients admitted to ED with discharge diagnosis of CAP | 28-day mortality |
| Kolditz (27), 2016 | 9327 | 63<sup>a</sup> | 5249 (56%) | Patients with CAP | 30-day mortality |
| LeGuen (28), 2017 | 182 | 72<sup>a</sup> | 88 (48%) | Patients reviewed by the RRT | 30-day mortality |
| Moskowitz (29), 2017 | 24164 | 64 | 12299 (51%) | Patients with suspected infection presented to ED | In-hospital mortality |
| Patidar (30), 2017 | 124 | 57 | NA | Cirrhotic patients hospitalized non-electively for infectious etiologies | 30-day mortality |
| Quinten (31), 2017 | 193 | 60 | 108 (56%) | Non-trauma patients in ED with suspected infection or sepsis | 28-day mortality |
| Ranzani (32), 2017 | 6874 | 66 | 4259 (62%) | Patients with clinical diagnosis of CAP | 30-day mortality |
| Rothman (33), 2017 | 3926 | NA | NA | Patients admitted to hospital with sepsis | In-hospital mortality |
| Seymour (5), 2016 | 66522 | 60 | 27446 (41%) | Patients with suspected infection | In-hospital mortality |
| Shetty (34),<br>2017 | 12555 | 50 <sup>a</sup> | 6585<br>(52%) | Patients with suspected infection, suspected or confirmed sepsis | Mortality and/or prolonged ICU stay $\geq 72$ hours |
| Singer (35),<br>2016 | 22530 | 54 | 10589<br>(47%) | ED patients whom qSOFA score could be calculated according to simultaneous reporting of vital signs and a MEWS score | In-hospital mortality |
| Szakmany<br>(36), 2017 | 380 | 74 <sup>a</sup> | 180<br>(47%) | Patients with high degree of clinical suspicion of infection | 30-day mortality |
| Tusgul (37),<br>2017 | 886 | 80 | 462<br>(52%) | Patients with suspected infection without alternative diagnosis, or microbiologically proven infection found in the ED workup | In-hospital mortality |
| Umemura<br>(38), 2017 | 387 | 74 <sup>a</sup> | 232<br>(60%) | ED patients admitted to ICU with diagnosis of severe sepsis | In-hospital mortality |
| Wang (39),<br>2016 | 477 | 73 <sup>a</sup> | 295<br>(62%) | Patients treated at ED with clinically diagnosed infection | 28-day mortality |
| Williams (7),<br>2017 | 8871 | 49 <sup>a</sup> | 4453<br>(50%) | ED patients admitted with a diagnosis indicating presumed or potential infection | 30-day mortality |
| **Long-term mortality** |  |  |  |  |  |
| Donnelly<br>(14), 2017 | 2593 | NA | NA | Admitted patients who meet the SIRS criteria, SOFA and qSOFA criteria | 1-year mortality |
| Quinten (31),<br>2017 | 193 | 60 | 108<br>(56%) | Non-trauma patients in ED with suspected infection or sepsis | 6-month mortality |
| Rannikko<br>(40), 2017 | 497 | 68 <sup>a</sup> | 262<br>(53%) | Adult patients admitted to the ED who had blood culture-positive sepsis | 90-day mortality |

Abbreviations: ED, emergency department; ICU, intensive care unit; ESS47, Emergency Symptoms and Signs algorithm for infection; CAP, community acquired pneumonia; NA, not available; SIRS, systemic inflammatory response syndrome; SOFA, Sequential organ failure assessment; qSOFA, quick sequential organ failure assessment; SK, “Sepsis Kills”; RRT, Rapid Response Team; MEWS, Modified Early Warning System.
<sup>a</sup> median

The prognostic accuracy of the qSOFA was evaluated in different countries, with most studies conducted in the United States of America and Europe, followed by Asia countries, Africa, New Zealand and Australia. Among all the articles included, six of them were conducted in Asia countries. The cut-off values of the Glasgow Coma Scale (GCS) used for altered mentation in the qSOFA included GCS less than 15, 14 and 13 respectively, and the remaining studies only stated altered mentation. A total of 35 studies reported on the prognostic accuracy of the qSOFA for short-term mortality^5–39^, while only 3 articles reported on long-term mortality ^14, 31, 40^.

### The qSOFA and short-term mortality

In this meta-analysis, 35 studies with 269,544 patients reported on the prognostic accuracy of the qSOFA and short-term mortality, including 27 retrospective studies ^5–8,10,11,13–16,18,20,22,24–27,29,32–35,37–40^, while 8 studies were prospective studies ^9,12,17,19,21,23,28,30,36^. Due to the heterogeneity of the inclusion criteria, a random-effects model was used to calculate the pooled sensitivity and specificity of the included studies. The forest plot for the sensitivity and specificity of the qSOFA predicting short-term mortality is shown in Figure 2. The pooled sensitivity was 48% and the specificity was 86%. The pooled odds ratio (OR) was 5.5 (95% CI: 5.32-5.72; Q =566.67; degree of freedom, df = 34; p<0.01), indicating that an elevated qSOFA score was associated with increased short-term mortality. The forest plot for the odds ratio is shown in Figure 3. We detected significant heterogeneity according to the heterogeneity tests (Tau = 24%; I^2^ = 94%; p<0.001). Publication bias was not detected as shown in the funnel plot (S1 Figure). Egger’s regression and Begg’s test revealed no statistical significance with p=0.84 (2-tailed) and p=0.46 respectively, indicating no publication bias (S8 Table).

**Figure 2.**
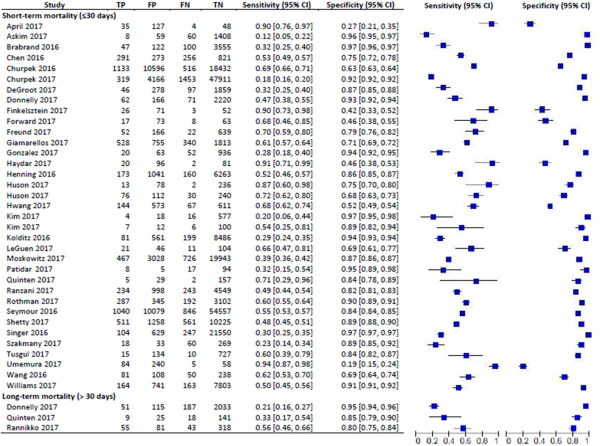
Sensitivity and Specificity of quick Sepsis-Related Organ Failure Assessment (qSOFA) in Predicting Short-term and Long-term Mortality. Studies included into the meta-analysis and their corresponding sensitivity and specificity of quick Sepsis-Related Organ Failure Assessment (qSOFA) values in predicting short- and long-term mortality is shown using a forest plot.

**Figure 3.**
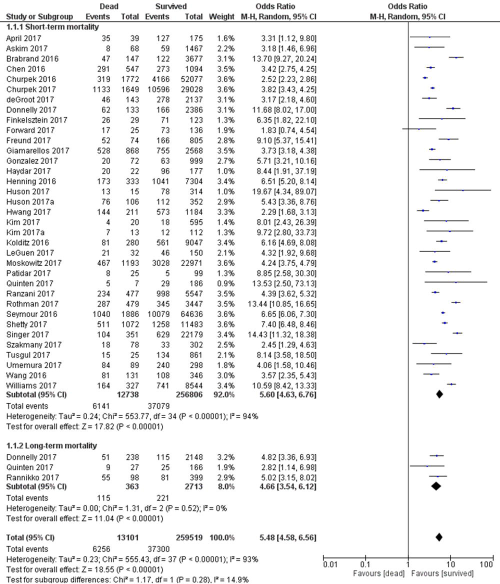
Odds Ratio of quick Sepsis-Related Organ Failure Assessment (qSOFA) in Predicting Short-term and Long-term Mortality. Odds of each study is shown in the forest plot. All studies found odds ratio of > 1 for quick Sepsis-Related Organ Failure Assessment (qSOFA) in predicting short- and long-term mortality.

### The qSOFA and long-term mortality

Only three studies with a total of 3,076 patients reported on the prognostic accuracy of the qSOFA and long-term mortality. Among these studies, two were retrospective ^14,40^ and one was prospective study ^31^. The forest plot for the sensitivity and specificity of the qSOFA for predicting long-term mortality is shown in Figure 2. The pooled sensitivity and specificity were calculated using a random-effects model, which yielded a pooled sensitivity of 32% and a pooled specificity of 92%. The three studies reported distinct mortality intervals, including 90-day mortality (sensitivity 56%, specificity 79%) ^40^, 6-month mortality (sensitivity 33%, specificity 85%) ^31^ and 12-month mortality (sensitivity 21%, specificity 95%) ^14^. The forest plot for the odds ratio is shown in Figure 3. The pooled OR was 4.7 (95% CI: 3.54-6.12; Q = 1.31; df = 2), and the studies were homogenous (Tau = 0%; I^2^<.001; p=0.52). However, publication bias was not assessed due to the small number of studies included in the long-term mortality analysis.

## DISCUSSION

This systematic review and meta-analysis revealed that most of the included studies suggested that a qSOFA score of ≥ 2 was able to predict short and long-term mortality. A total of 35 studies were reviewed, and the quality of the studies varied. Most of the studies had good quality according to QUADAS-2. Seven studies showed evidence of bias (S2 and S3 Tables). These seven studies had excluded many missing data and missing data analysis were not mentioned.

Our analysis revealed that qSOFA score exhibited fair sensitivity and specificity in predicting mortality. The pooled specificity of qSOFA in this study was higher compared to the SIRS (66%)^41^. According to our analysis, qSOFA can predict sepsis mortality, with the odds of 5.5 for the short-term mortality and 4.7 for the long-term mortality.

All 36 papers reporting on short-term mortality showed clinical, methodological and statistical heterogeneity. Factors that may have contributed to the high heterogeneity include the mean age (ranging from 54 to 84 years old), various clinical settings, variation in the timing of qSOFA scoring, and broad range of clinical diagnosis and criteria. This heterogeneity contributed to a lower pooled sensitivity of the qSOFA that may not represent the actual accuracy of the qSOFA. However, this finding was expected as the study population were diverse and multiple confounding factors were present. All studies showed positive direction in the forest plot reflecting a high pooled odds ratio. The funnel plot revealed no publication bias for the studies investigating qSOFA in predicting short-term mortality. Recently, three new publications reported on qSOFA short-term mortality prediction with similar findings^42–44^. Nevertheless, these studies did not perform further analysis on qSOFA long-term mortality prediction nor compared its prognostic accuracy with short-term mortality.

We found three studies which reported on qSOFA prognostication for long-term mortality. These studies showed clinical and methodological heterogeneity, but they were statistically homogenous. The performance of the qSOFA in long-term mortality prediction was more specific but less sensitive compared to its performance in short-term mortality. In studies reporting on long-term mortality, we found that the qSOFA was more specific but less sensitive. Since age is a confounding factor for mortality, we postulate that aging may contribute to this observation. In addition, longer mortality periods may result in the inclusion of deaths from extraneous factors. Further studies will be important to provide insight into this intriguing finding.

The qSOFA score consists of 3 criteria, and ranges from 0-3 points, with 1 point each for systolic hypotension (<100 mmHg), tachypnoea (>22/min), and altered mentation^5^. The three criteria of the qSOFA are much related to the pathophysiology of sepsis leading to organ injury. In sepsis, profound changes occur in the endothelium, causing widespread tissue oedema due to increased leukocyte adhesion, vasodilation, a shift to a pro-coagulant state and loss of barrier function, resulting in low blood pressure. Likewise, endothelial changes in severe sepsis causes increased permeability of lung capillaries, leading to accumulation of exudate fluid in the interstitial spaces of the lung, which floods into the alveoli and induces alveolar epithelial barrier dysfunction. These changes lead to acute respiratory distress and tachypnoea due to perfusion-ventilation mismatch, arterial hypoxemia and reduced lung compliance. Furthermore, systemic endothelial dysfunction affects the blood-brain barrier. Inflammatory cytokines and cells entered the brain, causing perivascular oedema, leukoencephalopathy, oxidative stress, and global neurotransmitter alteration, resulting in altered mental status ^45^. These complex changes in sepsis contributes to the high specificity of the qSOFA in detecting sepsis.

Sepsis was redefined in 2016 and the qSOFA was introduced to replace the SIRS as a better criteria for sepsis. The advantage of the qSOFA is that it can be repeatedly performed over time without laboratory investigations, which can be time-consuming ^25^. Since sepsis can deteriorate in a short period of time, a simple screening tool for early detection is warranted. The SIRS criteria introduced in previous sepsis definitions^3,46^ as a screening tool for sepsis was found to be overly sensitive relative to its specificity ^6^. A screening tool with high sensitivity and poor specificity can result in an excessive number of false positives, leading to unnecessary diagnostic or therapeutic procedures. Over-diagnosing patients poses a significant economic impact and further increases patients’ medical burden. In addition to qSOFA scoring, several publications have suggested lactate level could be of valuable biomarker when added to the original qSOFA score and may improve its prognostic value^21, 29, 34^. These studies provide insight into modification of the qSOFA which may improve its sensitivity and efficacy in detecting patients with sepsis. Efforts to modify the qSOFA could consider combining the present scoring criteria with other sepsis biomarkers such as C-reactive protein (CRP), procalcitonin (PCT), and serum secretory phospholipase A2-IIa (spla2-IIa)^46–49^.

Although the qSOFA exhibited high specificity and low sensitivity in most of the studies included in our meta-analysis, seven papers showed contradictory results. The studies reported that qSOFA was highly sensitive but had poor specificity. On further analysis, four of the studies had sample populations comprised of patients who were directly admitted from the ED to the ICU^8,15,24,38^, and two other studies included high numbers of Human Immunodeficiency Virus carriers ^22,23^. The remaining paper had a distinct study population including elderly and disabled patients, in whom assessment of altered mental status was regarded as challenging ^20^. The population included in these studies were more specific and likely to present to the ED with greater illness severity. Due to the specificity of these study populations, patients in these studies tended to be screened as positive as reflected by the identification of more true-positive patients compared to the other studies’ populations, resulting in heightened sensitivity of the qSOFA.

## LIMITATIONS

In this meta-analysis, we successfully retrieved all full-texts and a standardized tool was used to examine the quality of the included papers. One limitation of our analysis was the inclusion of too few articles reporting on long-term mortality. Second, we discovered that the study populations were substantially diverse, as some included specific groups of patients with infection. However, all of the included patients fulfilled our inclusion criterion of patients with suspected infection. Since random sampling was not performed in most of the studies included, a sampling bias is likely. Some studies had combined outcomes of mortality and/or ICU admission, thus complicating precise categorization of outcomes^10,34^. We classified in-hospital mortality as short-term mortality. Since in-hospital mortality may be longer than 30 days, this assumption may lead to a misclassification bias and mask the true predictive ability of the qSOFA. Most of the included studies were retrospective studies, posing a certain disadvantage as these studies relied on available medical records. Therefore, missing records or data may have influenced the results and the predictive accuracy of qSOFA in the current analysis. In addition, most of the studies were single-centered with variability across methods and study designs, which contributed to heterogeneity. Multiple confounders were likely to coexist, which may jeopardize the validity of these studies. Future research should consider prospective randomization in sampling method to minimize sampling bias. More studies exploring the qSOFA for long-term mortality prediction should be conducted in the near future.

## CONCLUSION

This meta-analysis revealed that the qSOFA score had a poor sensitivity but moderate specificity for both short and long-term mortality prediction in patients with suspected infection. Compared to short-term mortality, the qSOFA had superior predictive ability for long-term mortality. Further research on modification of qSOFA may improve its sensitivity in detecting patients with sepsis for prompt intervention. In general, qSOFA score may be a cost-effective tool for sepsis prognostication outside of the ICU setting.

**Supplementary Information** is available in the online version of the paper

**S1 Table. Table showing search strategy for both MEDLINE and SCOPUS including the keywords used and results from the database.**

**S2 Table. A summary of QUADAS-2 results for included studies from MEDLINE assessing the risk of bias and applicability concerns.** Three studies from MEDLINE showed evidence of bias.

**S3 Table. A summary of QUADAS-2 results for included studies from SCOPUS assessing the risk of bias and applicability concerns.** Four studies from SCOPUS showed evidence of bias.

**S4 Table. A detailed summary of type of study, subjects included, methodology and outcome of selected studies from MEDLINE**

**S5 Table. A detailed summary of type of study, subjects included, methodology and outcome of selected studies from SCOPUS.**

**S6 Table. Table showing list of articles excluded by title and abstract, including reason of exclusion.** A total of 41 studies were excluded by title and abstract.

**S7 Table. Table showing list of included and excluded full-text articles, and reason of exclusion.** A total of 36 studies were included after retrieving full-text articles and three studies were excluded due to incomplete data.

**S1 Figure. Funnel plot showing publication bias for short-term mortality.**

**S8 Table. Egger and Begg’s test were done for small-study effects, and no publication bias was detected.**

## Acknowledgement

The authors wish to thank Universiti Kebangsaan Malaysia for funding of the manuscript. This study was funded by grant Fundamental Research Grant Scheme (FRGS/1/2014/SKK01/UKM/03/3), Prototype Research Grant Scheme (PRGS/1/2017/STG05/UKM/03/1) from the Ministry of Higher Education, Malaysia and Fundamental Fund, FF-2018-015 from Faculty of Medicine, Universiti Kebangsaan Malaysia.

## Author Contribution

Dr Tan has full access to all the data in the study and takes responsibility for the integrity of the data and the accuracy of the data analysis.Study concept and design: Tan. Acquisition of data: Tan, Tang, Ching. Analysis and interpretation of data: All authors. Drafting of the manuscript: All authors. Critical revision of the manuscript for important intellectual content: All authors. Statistical analysis: Tan, Tang, Ching, Noraidulakma. Obtained funding: Tan

## Author Information

Reprints and permissions information is available at www.nature.com/reprints. The authors declare no competing interests. Correspondence and requests for materials should be addressed to.

## REFERENCES

1. Singer, M., et al. The Third International Consensus Definitions for Sepsis and Septic Shock (Sepsis-3). JAMA 315, 801–810 (2016).

2. Fleischmann, C., et al. Assessment of Global Incidence and Mortality of Hospital-treated Sepsis. Current Estimates and Limitations. American journal of respiratory and critical care medicine 193, 259–272 (2016).

3. Levy, M.M., et al. 2001 SCCM/ESICM/ACCP/ATS/SIS International Sepsis Definitions Conference. Intensive Care Medicine 29, 530–538 (2003).

4. Churpek, M.M., Zadravecz, F.J., Winslow, C., Howell, M.D. & Edelson, D.P. Incidence and Prognostic Value of the Systemic Inflammatory Response Syndrome and Organ Dysfunctions in Ward Patients. American journal of respiratory and critical care medicine 192, 958–964 (2015).

5. Seymour, C.W., et al. Assessment of Clinical Criteria for Sepsis: For the Third International Consensus Definitions for Sepsis and Septic Shock (Sepsis-3).[Erratum appears in JAMA. 2016 May 24-31;315(20):2237; PMID: 27218643]. JAMA 315, 762–774 (2016).

6. Churpek, M.M., et al. Quick Sepsis-related Organ Failure Assessment, Systemic Inflammatory Response Syndrome, and Early Warning Scores for Detecting Clinical Deterioration in Infected Patients outside the Intensive Care Unit. American Journal of Respiratory & Critical Care Medicine 195, 906–911 (2017).

7. Williams, J.M., et al. Systemic Inflammatory Response Syndrome, Quick Sequential Organ Function Assessment, and Organ Dysfunction: Insights From a Prospective Database of ED Patients With Infection. Chest 151, 586–596 (2017).

8. April, M.D., et al. Sepsis Clinical Criteria in Emergency Department Patients Admitted to an Intensive Care Unit: An External Validation Study of Quick Sequential Organ Failure Assessment. Journal of Emergency Medicine 52, 622–631 (2017).

9. Askim, Å., et al. Poor performance of quick-SOFA (qSOFA) score in predicting severe sepsis and mortality - a prospective study of patients admitted with infection to the emergency department. Scandinavian Journal of Trauma, Resuscitation and Emergency Medicine 25 (2017).

10. Brabrand, M., Havshoj, U. & Graham, C.A. Validation of the qSOFA score for identification of septic patients: A retrospective study. European Journal of Internal Medicine 36, e35–e36 (2016).

11. Chen, Y.X., Wang, J.Y. & Guo, S.B. Use of CRB-65 and quick Sepsis-related Organ Failure Assessment to predict site of care and mortality in pneumonia patients in the emergency department: a retrospective study. Critical Care (London, England) 20, 167 (2016).

12. Churpek, M.M., Snyder, A., Sokol, S., Pettit, N.N. & Edelson, D.P. Investigating the Impact of Different Suspicion of Infection Criteria on the Accuracy of Quick Sepsis-Related Organ Failure Assessment, Systemic Inflammatory Response Syndrome, and Early Warning Scores. Critical Care Medicine (2017).

13. de Groot, B., et al. The most commonly used disease severity scores are inappropriate for risk stratification of older emergency department sepsis patients: an observational multicentre study. Scandinavian Journal of Trauma, Resuscitation and Emergency Medicine 25, 91 (2017).

14. Donnelly, J.P., Safford, M.M., Shapiro, N.I., Baddley, J.W. & Wang, H.E. Application of the Third International Consensus Definitions for Sepsis (Sepsis-3) Classification: a retrospective population-based cohort study. The Lancet Infectious Diseases 17, 661–670 (2017).

15. Finkelsztein, E.J., et al. Comparison of qSOFA and SIRS for predicting adverse outcomes of patients with suspicion of sepsis outside the intensive care unit. Critical Care 21 (2017).

16. Forward, E., et al. Predictive validity of the qSOFA criteria for sepsis in non-ICU inpatients. Intensive Care Medicine 43, 945–946 (2017).

17. Freund, Y., et al. Prognostic Accuracy of Sepsis-3 Criteria for In-Hospital Mortality Among Patients With Suspected Infection Presenting to the Emergency Department. JAMA 317, 301–308 (2017).

18. Giamarellos-Bourboulis, E.J., et al. Validation of the new Sepsis-3 definitions: proposal for improvement in early risk identification. Clinical Microbiology and Infection 23, 104–109 (2017).

19. González Del Castillo, J., et al. Prognostic accuracy of SIRS criteria, qSOFA score and GYM score for 30-day-mortality in older non-severely dependent infected patients attended in the emergency department. European Journal of Clinical Microbiology and Infectious Diseases, 1–9 (2017).

20. Haydar, S., Spanier, M., Weems, P., Wood, S. & Strout, T. Comparison of qSOFA score and SIRS criteria as screening mechanisms for emergency department sepsis. American Journal of Emergency Medicine (2017).

21. Henning, D.J., et al. An Emergency Department Validation of the SEP-3 Sepsis and Septic Shock Definitions and Comparison With 1992 Consensus Definitions. Annals of Emergency Medicine (2016).

22. Huson, M.A.M., Kalkman, R., Grobusch, M.P. & van der Poll, T. Predictive value of the qSOFA score in patients with suspected infection in a resource limited setting in Gabon. Travel Medicine and Infectious Disease 15, 76–77 (2017).

23. Huson, M.A.M., et al. Application of the qSOFA score to predict mortality in patients with suspected infection in a resource-limited setting in Malawi. Infection, 1–4 (2017).

24. Hwang, S.Y., et al. Low Accuracy of Positive qSOFA Criteria for Predicting 28-Day Mortality in Critically Ill Septic Patients During the Early Period After Emergency Department Presentation. Annals of Emergency Medicine (2017).

25. Kim, M., et al. Predictive performance of the quick Sequential Organ Failure Assessment score as a screening tool for sepsis, mortality, and intensive care unit admission in patients with febrile neutropenia. Supportive Care in Cancer 25, 1557–1562 (2017).

26. Kim, M.W., Lim, J.Y. & Oh, S.H. Mortality prediction using serum biomarkers and various clinical risk scales in community-acquired pneumonia. Scandinavian Journal of Clinical and Laboratory Investigation 77, 486–492 (2017).

27. Kolditz, M., et al. Vergleich der qSOFA- und CRB-Kriterien zur Risikoprädiktion bei Patienten mit CAP: erste multizentrische Validierung des qSOFA bei CAP. Pneumologie (Stuttgart, Germany) 70, 826–830 (2016).

28. LeGuen, M., et al. Frequency and significance of qSOFA criteria during adult rapid response team reviews: A prospective cohort study. Resuscitation 122, 13–18 (2018).

29. Moskowitz, A., et al. Quick Sequential Organ Failure Assessment and Systemic Inflammatory Response Syndrome Criteria as Predictors of Critical Care Intervention Among Patients With Suspected Infection. Critical care medicine 45, 1813–1819 (2017).

30. Patidar, K.R., et al. No Association Between Quick Sequential Organ Failure Assessment and Outcomes of Patients With Cirrhosis and Infections. Clinical Gastroenterology and Hepatology 15, 1803–1804 (2017).

31. Quinten, V.M., van Meurs, M., Wolffensperger, A.E., ter Maaten, J.C. & Ligtenberg, J.J.M. Sepsis patients in the emergency department: stratification using the Clinical Impression Score, Predisposition, Infection, Response and Organ dysfunction score or quick Sequential Organ Failure Assessment score? European Journal of Emergency Medicine (2017).

32. Ranzani, O.T., et al. New Sepsis Definition (Sepsis-3) and Community-acquired Pneumonia Mortality. A Validation and Clinical Decision-Making Study. American journal of respiratory and critical care medicine 196, 1287–1297 (2017).

33. Rothman, M., et al. Sepsis as 2 problems: Identifying sepsis at admission and predicting onset in the hospital using an electronic medical record–based acuity score. Journal of Critical Care 38, 237–244 (2017).

34. Shetty, A., et al. Lactate ≥2 mmol/L plus qSOFA improves utility over qSOFA alone in emergency department patients presenting with suspected sepsis. EMA - Emergency Medicine Australasia 29, 626–634 (2017).

35. Singer, A.J., Ng, J., Thode, H.C., Jr., Spiegel, R. & Weingart, S. Quick SOFA Scores Predict Mortality in Adult Emergency Department Patients With and Without Suspected Infection. Annals of Emergency Medicine 69, 475–479 (2017).

36. Szakmany, T., et al. Defining sepsis on the wards: Results of a multi-centre point-prevalence study comparing two sepsis definitions. Anaesthesia (2017).

37. Tusgul, S., Carron, P.N., Yersin, B., Calandra, T. & Dami, F. Low sensitivity of qSOFA, SIRS criteria and sepsis definition to identify infected patients at risk of complication in the prehospital setting and at the emergency department triage. Scandinavian Journal of Trauma, Resuscitation and Emergency Medicine 25 (2017).

38. Umemura, Y., et al. Assessment of mortality by qSOFA in patients with sepsis outside ICU: A post hoc subgroup analysis by the Japanese Association for Acute Medicine Sepsis Registry Study Group. Journal of Infection and Chemotherapy (2017).

39. Wang, J.Y., Chen, Y.X., Guo, S.B., Mei, X. & Yang, P. Predictive performance of quick Sepsis-related Organ Failure Assessment for mortality and ICU admission in patients with infection at the ED. American Journal of Emergency Medicine 34, 1788–1793 (2016).

40. Rannikko, J., Syrjänen, J., Seiskari, T., Aittoniemi, J. & Huttunen, R. Sepsis-related mortality in 497 cases with blood culture-positive sepsis in an emergency department. International Journal of Infectious Diseases 58, 52–57 (2017).

41. Hoeboer, S.H., van der Geest, P.J., Nieboer, D. & Groeneveld, A.B. The diagnostic accuracy of procalcitonin for bacteraemia: a systematic review and meta-analysis. Clinical microbiology and infection: the official publication of the European Society of Clinical Microbiology and Infectious Diseases 21, 474–481 (2015).

42. Fernando, S.M., et al. Prognostic Accuracy of the Quick Sequential Organ Failure Assessment for Mortality in Patients With Suspected Infection. Ann Intern Med 168, 266–275 (2018).

43. Song, J.-U., Sin, C.K., Park, H.K., Shim, S.R. & Lee, J. Performance of the quick Sequential (sepsis-related) Organ Failure Assessment score as a prognostic tool in infected patients outside the intensive care unit: a systematic review and meta-analysis. Critical Care 22, 28 (2018).

44. Maitra, S., Som, A. & Bhattacharjee, S. Accuracy of quick Sequential Organ Failure Assessment (qSOFA) score and systemic inflammatory response syndrome (SIRS) criteria for predicting mortality in hospitalized patients with suspected infection: A meta-analysis of observational studies: Predictive accuracy of qSOFA: A meta-analysis. Clinical Microbiology and Infection (2018).

45. Gotts, J.E. & Matthay, M.A. Sepsis: pathophysiology and clinical management. BMJ 353 (2016).

46. Bone, R.C., et al. Definitions for sepsis and organ failure and guidelines for the use of innovative therapies in sepsis. The ACCP/SCCM Consensus Conference Committee. American College of Chest Physicians/Society of Critical Care Medicine. Chest 101, 1644–1655 (1992).

47. Vijayan, A.L., et al. Procalcitonin: a promising diagnostic marker for sepsis and antibiotic therapy. Journal of Intensive Care 5, 51 (2017).

48. Tan, T.L. & Goh, Y.Y. The role of group IIA secretory phospholipase A2 (sPLA2-IIA) as a biomarker for the diagnosis of sepsis and bacterial infection in adults—A systematic review. PLOS ONE 12, e0180554 (2017).

49. Faix, J.D. Biomarkers of sepsis. Critical Reviews in Clinical Laboratory Sciences 50, 23–36 (2013).

